# Homodimerization of CB_2_ cannabinoid receptor triggered by a bivalent ligand enhances cellular signaling

**DOI:** 10.1101/2024.05.10.593612

**Authors:** Gemma Navarro, Marc Gómez-Autet, Paula Morales, Joan Biel Rebassa, Claudia Llinas del Torrent, Nadine Jagerovic, Leonardo Pardo, Rafael Franco

**Author notes:** These authors contributed equally to this work (Gemma Navarro, Marc Gómez-Autet, Paula Morales).

## Abstract

G protein-coupled receptors (GPCRs) exist within a landscape of interconvertible conformational states and in dynamic equilibrium between monomers and higher-order oligomers, both influenced by ligand binding. Here, we have shown that a homobivalent ligand formed by equal chromenopyrazole moieties as pharmacophores, connected by 14 methylene units, can modulate the dynamics of the cannabinoid CB_2_ receptor (CB_2_R) homodimerization by simultaneously binding both protomers of the CB_2_R-CB_2_R homodimer. Computational and pharmacological experimentals showed that one of the ligand pharmacophores binds to the orthosteric site of one protomer, and the other pharmacophore to a membrane-oriented pocket between transmembranes 1 and 7 of the partner protomer. This provides unique pharmacological properties, such as increased potency in G_i_ binding and increased recruitment of β-arrestin. Thus, by modulating dimerization dynamics, it may be possible to fine-tune CB_2_R activity with potentially improved therapeutic outcomes.

**HIGHLIGHTS:**

- A homobivalent ligand of CB_2_R (PM369) modulates the dynamics of receptor homodimerization
- PM369 binds to the orthosteric site of one protomer and to a complementary, membrane-facing, site of the other protomer
- PM369 triggers CB_2_R homodimerization via the TM 1/7 interface that provides unique pharmacological properties
- PM369 potentiates signaling, increased potency in G_i_ binding and increased recruitment of β-arrestin
- These results highlight new approaches to control GPCR signaling

**GRAPHICAL ABSTRACT:** 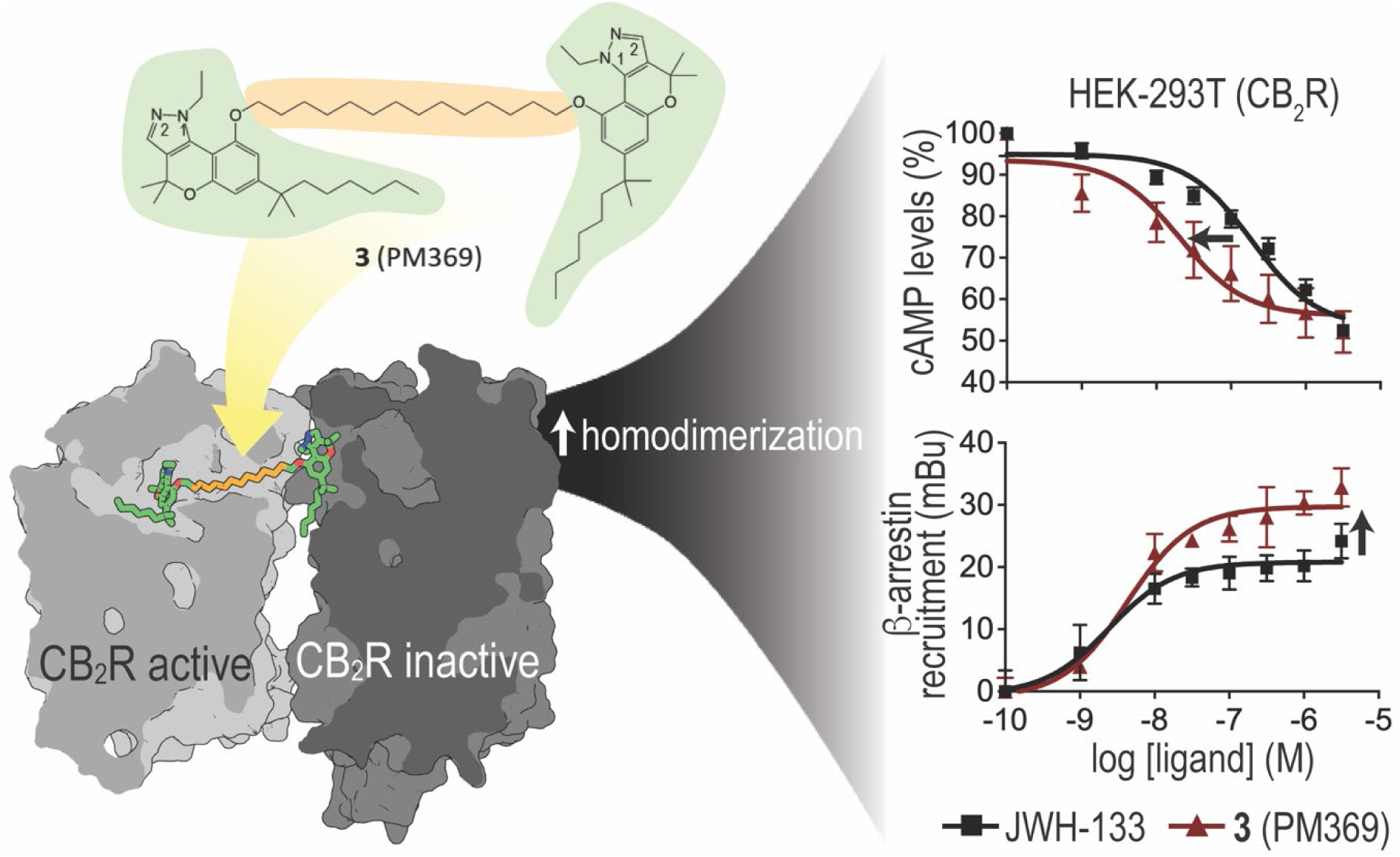

## 1. Introduction

In class A G protein-coupled receptors (GPCRs), endogenous ligands bind to a conserved pocket within the seven transmembrane helices (TMs) of the receptor that optimally accommodates their electrostatic and steric properties [1]. Agonist binding to this extracellular orthosteric site triggers local structural changes, which vary among GPCR families, in the immediate proximity of the highly conserved PIF motif. The agonist-induced arrangement of the PIF side chains is transmitted into larger-scale helix movements at the intracellular site via the highly conserved NPxxY and DRY motifs and Y5.58 [2,3], so that an intracellular cavity is opened through the outward movement of TMs 5 and 6 [4]. Opening of this cavity permits the C-terminal α5 helix of G-proteins [5], the finger loop of β-arrestins [6], or the N-terminal αN helix of GPCR kinases [7] to bind the receptor for downstream signaling pathways. The greater efficacy of some exogenous ligands to stimulate a particular transducer is known as ligand bias [8].

The size of a G protein or β-arrestin is larger than the 7TM domain of a GPCR and, despite a monomeric GPCR can activate a G protein [9], it is also feasible a 2:1 (receptor:G protein/arrestin) stoichiometry [10]. Single-molecule microscopy techniques [11] have shown that GPCRs are present in a dynamic equilibrium between monomers, dimers, and tetramers [12–15]. We have described that the formation of homo- or hetero-oligomers [16–21] may alter the signaling response compared to receptors expressed individually on the cell surface [16,17,21]. In fact, the formation of oligomers can influence G protein or β-arrestin binding [15,22]. However, the mechanism by which the second protomer of the homodimer modulates the G protein or β-arrestin remains unknown.

The recent analysis of the pocketome of GPCRs has shown multitude non-conserved pockets in addition to the orthosteric pocket [23]. These are located at the membrane-facing part of the receptor, at the entrance (vestibule, secondary, or metastable site or exosite) of the orthosteric site, as previously suggested by molecular dynamics (MD) simulations [24–26]. Pharmacophoric moieties that bind at the orthosteric site exhibit high-affinity for the receptor, whereas pharmacophores that bind at these non-conserved pockets confer receptor selectivity [27–29]. The bivalent approach [30] bridges these two pharmacophores covalently by a spacer for selectivity, off-rates, and signaling bias [31–33].

In GPCRs activated by signaling molecules derived from lipid species, such as cannabinoid, sphingosine-1-phosphate, and lysophosphatidic acid, the extracellular N-terminus and extracellular loop 2 fold over the ligand binding pocket [25]. Thus, the entrance of ligands to the orthosteric site located within the 7TM domain occurs through the membrane bilayer [34–36]. This is the case of cannabinoid receptors, type 1 (CB_1_R) and type 2 (CB_2_R), that respond to endocannabinoids with long hydrophobic tails such as anandamide and 2-arachidonoyl glycerol, as well as phytocannabinoids like Δ⁹-tetrahydrocannabinol [37,38]. The pathway of ligand entry to CB_2_R, determined by MD simulations, defines two transient binding sites [39]: a membrane-facing pocket between TMs 1 and 7 [40] and a bundle-facing allosteric pocket located near the orthosteric site [41]. In a previous study, we reported the characterization of the first bitopic homobivalent ligand for the CB_2_R (**4**; Figure 1) [40]. Combining pharmacological experiments, MD simulations, and site-directed mutagenesis, we showed that the symmetrical pharmacophore units of **4** simultaneously bind the orthosteric site and the membrane-facing pocket. Here, we report that a homobivalent ligand with a precise linker size can modulate the dynamics of CB_2_R homodimerization, as it has been suggested for other bivalent compounds (e.g. for dopamine D_2_-D_2_ receptor homodimer [42] and for adenosine A_2A_-dopamine D_2_ receptor heteromer [43]). Importantly, we show that the formation of CB_2_R homodimerization, through a specific TM interface, promoted by the bivalent ligand provides unique pharmacological properties, such as increased potency in G_i_ binding and enhanced β-arrestin recruitment. Thus, the research described herein highlights the intricacies of GPCR modulation and presents a novel approach, using homobivalent ligands to influence the dynamics of GPCR homodimerization and signaling.

**Figure 1.**
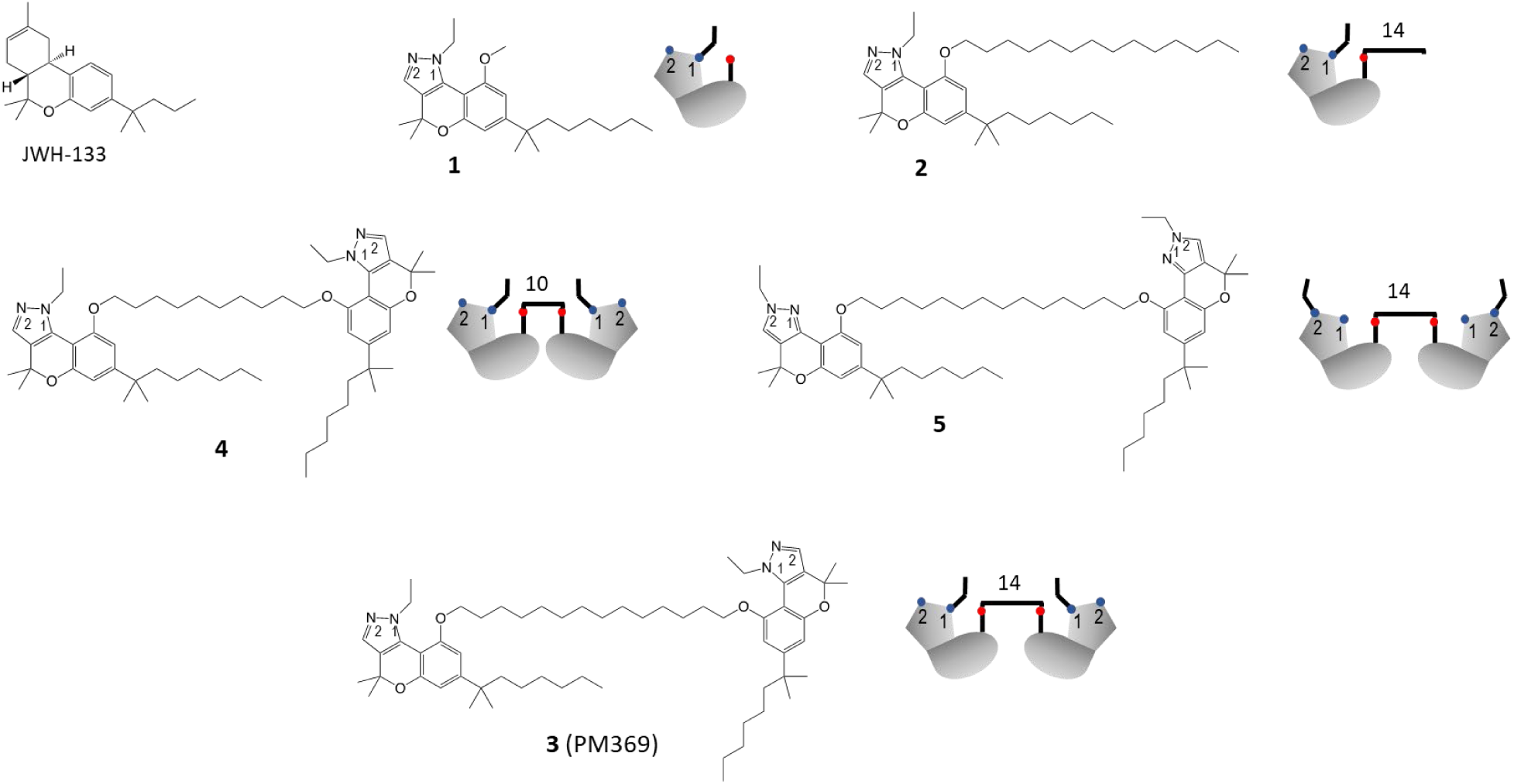
Chemical structures and cartoon representations of compounds **3** (PM369), **1**, **2**, **4**, and **5**, previously reported by us [40], and the reference CB_2_R agonist JWH-133, used in this manuscript.

## 2. Material and Methods

### 2.1. Chemicals

The synthesis of compounds **1**-**2**, **3** (PM369), **4**-**5**, have been previously described by us [40,44]. The reference CB_2_R agonist JWH-133 (Dimethylbutyl-deoxy-Δ^8^-tetrahydrocannabinol; CAS 259869-55-1) has been purchased to Tocris Bioscience (Bristol, UK).

### 2.2. Cell Culture and Transient Transfection

HEK-293T cells were grown in 6-well plates in Dulbecco’s modified Eagle’s medium (DMEM) (15-013-CV, Corning, NY, USA) supplemented with 2 mM L-glutamine, 100 U/ml penicillin/streptomycin, MEM Non-Essential Amino Acids Solution (1/100) and 5% (v/v) heat inactivated Fetal Bovine Serum (FBS) (all supplements were from Invitrogen, (Paisley, Scotland, UK)). Cells were cultured in a humid atmosphere of 5% CO_2_ at 37°C. After 24 h in culture, cells were transiently transfected with the corresponding plasmids by the polyethyleneimine (PEI) method. PEI is a commonly used cationic polymer that form complexes with negatively charged nucleic acids and facilitate their uptake into cells. The corresponding cDNAs diluted in 150 mM NaCl were mixed with PEI (5.5 mM in nitrogen residues, Sigma-Aldrich, St. Louis, MO, USA) also prepared in 150 mM NaCl for 10 min. The PEI-DNA complex mixture was added to cells dropwise in non-supplemented DMEM, and the plate was swirled gently to ensure even distribution. Cells were incubated at 37°C in a humid atmosphere of 5% CO_2_ for 4–6 h. After incubation, the transfection medium was removed and replaced with complete culture medium, and assays were performed after 48 h incubation at 37°C in a 5% CO_2_ humid atmosphere.

### 2.3 Bioluminescence Resonance Energy Transfer (BRET^2^)

HEK-293T cells growing in 6-well plates were transiently co-transfected with a constant amount of cDNA encoding for human CB_2_R fused to Renilla luciferase (CB_2_R-Rluc) and with increasing amounts of cDNAs corresponding to CB_2_R fused to the green fluorescent protein GFP^2^ (CB_2_R-YFP). The negative controls were performed uing gamma-amino butyric acid receptor 1B (GABA_1B_R) fused to Rluc. 48 h post-transfection, cells were washed twice in quick succession in HBSS (137 mM NaCl; 5 mM KCl; 0.34 mM Na_2_HPO4; 0.44 mM KH_2_PO_4_; 1.26 mM CaCl_2_; 0.4 mM MgSO_4_; 0.5 mM MgCl_2_ and 10 mM HEPES, pH 7.4) supplemented with 0.1% glucose (w/v), detached by gently pipetting and resuspended in the same buffer. To assess the number of cells per plate, we determined protein concentration using a Bradford assay kit (Bio-Rad, Munich, Germany) with bovine serum albumin dilutions as standards, adjusting cells to a concentration of 0.2 mg/mL. To quantify GFP^2^-fluorescence expression, we distributed the cells (20 μg protein) in 96-well microplates (black plates with a transparent bottom; Porvair, Leatherhead, UK). Fluorescence was read using a Mithras LB 940 (Berthold, Bad Wildbad, Germany) equipped with a high-energy xenon flash lamp, using a 10-nm bandwidth excitation and emission filters at 410 and 510 nm, respectively. GFP^2^-fluorescence expression was determined as the fluorescence of the sample minus the fluorescence of cells expressing protein-Rluc alone. For the BRET measurements, the equivalent of 20 μg of cell suspension was distributed in 96-well microplates (white plates; Porvair), and we added 5 μM Deep Blue C (PJK GMBH, Kleinblittersdorf, Germany). Then, 30 seconds after Deep Blue C addition, the readings were collected using a Mithras LB 940 (Berthold, Bad Wildbad, Germany), which allowed the integration of the signals detected in the short-wavelength filter at 410 nm (400–430 nm) and the long-wavelength filter at 510 nm (500–530 nm). To quantify receptor-Rluc expression, we performed luminescence readings 10 min after addition of 5 μM coelenterazine H. The net BRET is defined as [(long-wavelength emission)/ (short-wavelength emission)]-Cf where Cf corresponds to [(long-wavelength emission)/ (short-wavelength emission)] for the Rluc construct expressed alone in the same experiment. The BRET curves were fitted assuming a single phase by a non-linear regression equation using the GraphPad Prism software (San Diego, CA, USA). BRET values are given as milli BRET units (mBU: 1000 × net BRET).

### 2.4. Preparation of plasmids coding for CB_2_R containing point mutations

Mutations were produced using the QuikChange® Site-Directed Mutagenesis Kit. The cDNA for hCB_2_R, cloned into pcDNA3.1, was amplified using sense and antisense primers harboring the triplets for the desired point mutation (Pfu turbo polymerase was used). The nonmutated DNA template was digested for 1 h with DpnI. PCR products were used to transform XL1-blue supercompetent cells. Finally, positive colonies were tested by sequencing to select those expressing the correct DNA sequence.

### 2.5. ß-Arrestin 2 recruitment

HEK-293T cells were transiently co-transfected with cDNA coding for ß-arrestin 2-Rluc and with cDNA coding for CB_2_R-YFP. BRET experiments were performed 48 h after transfection. Cells were detached using HBSS containing 0.1% glucose, centrifuged for 5 min at 3,200 rpm and resuspended in the same buffer. Protein concentration was quantified by using the Bradford assay kit (Bio-Rad, Munich, Germany) and adjusted to 0,2 mg/mL. Hereafter, YFP fluorescence was quantified at 530 nm in a FluoStar Optima Fluorimeter (BMG Labtech, Offenburg, Germany) to quantify receptor-YFP expression upon excitation at 488 nm. To measure β-arrestin 2 recruitment, cells (20 µg of protein) were distributed in 96-well microplates (Corning 3600, white plates with white bottom) and were incubated for 10 min with antagonists. Cells were then stimulated with agonists prior to the addition of 5 µM coelenterazine H (Molecular Probes, Eugene, OR). BRET between β-arrestin 2-Rluc and receptor-YFP was determined and quantified 5 min after adding coelenterazine H. The readings were collected using a Mithras LB 940 (Berthold Technologies, Bad Wildbad, Germany), which allows the integration of the signals detected in the short-wavelength filter (485 nm) and the long wavelength filter (530 nm). To quantify protein-Rluc expression, luminescence readings were also collected 10 min after the addition of 5 µM coelenterazine H.

### 2.6. cAMP level determination

As the CB_2_R receptor couples to heteromeric G_i_ proteins, its activation by agonists leads to the inhibition of adenylate cyclase and the subsequent decrease of intracellular concentration of adenosine cyclic 3’,5’-monophosphate (cAMP). The concentration of this first messenger was determined using the Lance Ultra cAMP kit (Perkin Elmer, Waltham, MA, US). The method consists of a time-resolved fluorescence resonance energy transfer (TR-FRET) immunoassay in which endogenous cAMP competes with europium (Eu) chelate-labeled cAMP tracer for binding sites on a cAMP-specific antibody labeled with the ULight^TM^ dye. cAMP concentrations per cell or per mg protein were determined using a standard curve using pure unlabeled cAMP.

HEK-293T cells growing in 6-well plates were transiently transfected with the cDNA for the non-mutated or mutated human CB_2_R. 48 h post-transfection, medium was replaced by serum-free DMEM. Two hours later, cells were detached, isolated by centrifugation (5 min at 1500 rpm) and resuspended in ^cAMP^medium, which consists of DMEM containing 5 mM HEPES (pH 7.4), BSA (0,1 %) and 50 µM zardaverine (phosphodiesterase inhibitor that prevents degradation of cAMP). Determination was performed in 384-well plates (Perkin Elmer) using 4000 cells/well. Cells were incubated for 15 min with 2 µl of ligands prepared in ^AMP^medium or vehicle. Finally, cells were treated for 15 min with forskolin (500 nM). 15 min later cAMP-Europium (cAMP-Eu) (5 µl) and fluorophore-containing ULight^TM^ antibody (5 µl) were added. Incubation was prolonged for 1 h at 25°C and the PHERAstar Flagship reader equipped with an HTRF optical module (BMG Lab technologies, Offenburg, Germany) was used for measuring the 665/620 nm ratio.

### 2.7. In Situ Proximity Ligation Assay (PLA)

The proximity ligation assay (PLA) allows the detection of molecular interactions between two proteins in homologous cells and in fixed tissue sections. The presence/absence of receptor–receptor molecular interactions in the samples was detected using the Duolink II in Situ PLA Detection Kit (developed by Olink Bioscience, Uppsala, Sweden; and now distributed by Sigma-Aldrich as Duolink^®^, using PLA^®^ Technology).

HEK-293T cells were transfected with cDNAs for CB_2_R-Rluc and CB_2_R-GFP^2^ (5 µg cDNA each). Samples were treated with specific antibodies against Rluc (1/100; MAB4400, Millipore) and against GFP (1/100; A10262, Invitrogen) and processed using the PLA probes Duolink II antimouse plus and anti-rabbit minus).

Samples were fixed in 4% paraformaldehyde for 15 min and then washed twice with PBS containing 20 mM glycine before permeabilization with PBS-glycine containing 0.2% Triton X-100. After permeabilization, the samples were washed in PBS at room temperature and incubated in a preheated humidity chamber for 1 h at 37°C with the blocking solution provided in the PLA kit. Then, the samples were incubated overnight with the primary antibodies. Day after, samples were incubated for 1 h with PLA probe-linked secondary antibodies (1:100 dilutions for all antibodies) at 4 °C. After washing, the samples were incubated with the ligation solution for 1 h, and then washed and subsequently incubated with the amplification solution for 100 min (both steps at 37°C in a humid chamber). Nuclei were stained with Hoechst (1/100 from stock 1 mg/mL; Thermo Fisher). The samples were washed several times and mounted on glass slides with ShandonTM Immu-MountTM (9990402; ThermoFisher). Samples were observed under a Zeiss 880 confocal microscope (Carl Zeiss, Oberkochen, Germany) equipped with an apochromatic 63× oil-immersion objective (N.A. 1.4), and with 405 nm and 561 nm laser lines. For each field of view, a stack of two channels (one per staining) and 4 Z planes with a step size of 0.2 µm were acquired. The ratio r (number of red spots/cell) was determined on the maximum projection of each image stack using the Duolink Image tool software.

### 2.8. Computational methods

The GPCRdb refined structure [45] of inactive CB_2_R bound to the AM10257 antagonist (PDB id 5ZTY) [46] and the active CB_2_R-G_i_ complex bound to the WIN 55,212-2 agonists (6PT0) [47] were used. Fusion proteins and antibodies were removed, and stabilizing mutations were mutated to the native sequence. The CB_2_R-CB_2_R homodimer was built from the TM 1/7 dimeric interface observed in the crystal structure of the serotonin 5-HT_2C_ receptor (6BQH) [48]. The chromenopyrazole pharmacophore was modeled into the orthosteric binding cavity of active CB_2_R, using the structure of CB_2_R-G_i_ bound to HU308 agonist (PDB id 8GUS) [49] as a template; and docked into the membrane-facing pocket between TMs 1 and 7 of inactive CB_2_R using the Molecular Operating Environment (MOE) software (Chemical Computing Group Inc., Montreal, Quebec, Canada). The 14 methylene units of the spacer of homobivalent ligand **3** (PM369) were modelled connecting both pharmacophores through the tunnel between TMs 1 and 7 of active CB_2_R. The binding of β-arrestin to CB_2_R was modeled using the structure of β-arrestin bound to the CB_1_R (8WU1) [50] or (8WRZ) [51] as templates. These structures of the CB_2_R-CB_2_R homodimer were embedded in a lipid bilayer box, constructed using PACKMOL-memgen [52], containing 1-palmitoyl-2-oleoyl-sn-glycero-3-phosphocholine, water molecules, and monoatomic Na^+^ and Cl^-^ ions. MD simulations of these systems were performed with GROMACS 2019 using the protocol previously reported [53]. The analysis of the trajectories was performed using MDAnalysis [54] and GetContacts [55].

## 3. Results and Discussion

### 3.1. A homobivalent ligand of CB_2_R modulates the dynamics of receptor homodimerization

We have previously reported a series of symmetrical homobivalent ligands formed by equal chromenopyrazole moieties [44] as pharmacophores connected by spacers whose lengths vary from six to sixteen methylene units [40]. Among these compounds, there were two isomers that differed in the position of the N-ethyl in the pyrazole (N1- or N2-ethyl). Homobivalent N1-ethyl ligands with spacers lengths of 10, 12, or 14 methylene units and N2-ethyl ligands with 12 or 14 methylenes showed the best binding affinity for CB_2_R [40]. Computational models and mutagenesis experiments indicated that these linker lengths allow both pharmacophores at the ligand ends to bind simultaneously to the orthosteric site and to a membrane-facing site between TMs 1 and 7 [40]. From these models, we observed that one of the chromenopyrazole pharmacophores was located at the membrane-facing part, thus exposed to a putative second protomer. Taking into account this observation, we now test the ability of these compounds to modulate the dynamics of CB_2_R homodimerization. With this aim, we used bioluminescence resonance energy transfer (BRET^2^) assays in cells expressing CB_2_R fused with Renilla luciferase (CB_2_R-Rluc) as BRET donor and fused to the GFP^2^ green fluorescent protein (CB_2_R-GFP^2^) as BRET acceptor (see Methods). Homodimers of CB_2_R were detected in HEK-293T cells, whereas GABA_1B_R-Rluc, used as a negative control, could not interact with CB_2_R-GFP^2^ (Figure 2A). Remarkably, the BRET^2^_max_ value (284.6 ± 24.7 mBU) obtained with the N1-ethyl homobivalent ligand with 14 methylene units **3** (PM369) was markedly higher than those obtained in untreated cells (72.6 ± 4.3 mBU), or in the presence of the pharmacophore unit **1** (60.5 ± 4.2 mBU), the monovalent compound that lacks the second pharmacophore **2** (94.6 ±18.5 mBU), or a shorter N1-ethyl homobivalent ligand with 10 methylene units **4** (74.1 ± 15.3 mBU) (Figures 2C-2D). In summary, the increase BRET^2^_max_, which reflects greater homodimerization, requires a specific linker size of the homobivalent ligand that supposedly permits the interaction with the second CB_2_R protomer. It is also noteworthy that the homobivalent N2-ethyl ligand with 14 methylene units **5** did not increase BRET^2^_max_ compared to untreated cells (Figure 2B). This suggests that the pyrazole ring of the second pharmacophore plays a key role in the recognition of the second protomer of the CB_2_R homodimer.

**Figure 2.**
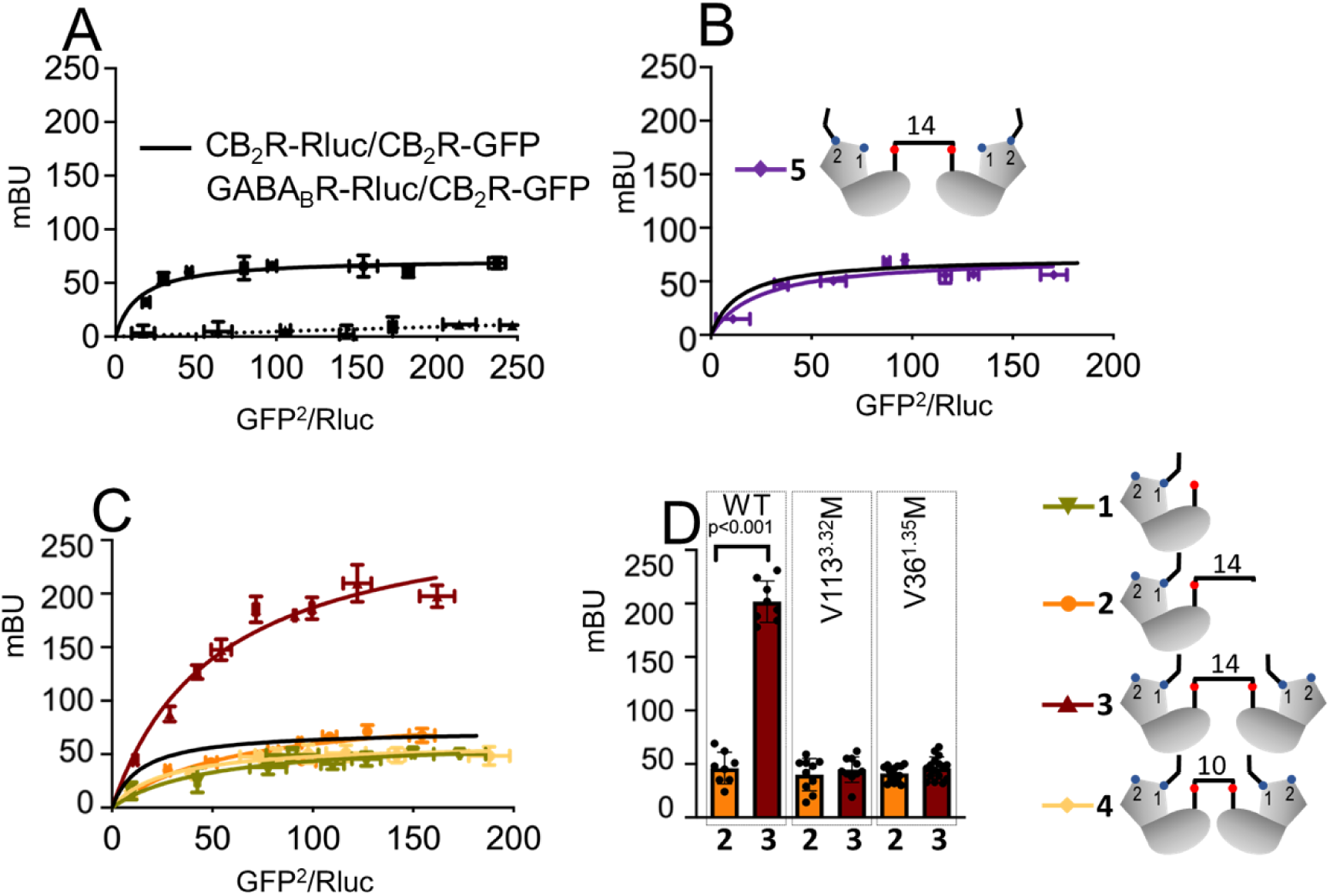
Homodimerization of CB_2_R in the presence of symmetrical homobivalent ligands. BRET assays were performed in HEK-293T cells transfected with a constant amount of cDNA for CB_2_R-Rluc (0.3 μg) (A-C) or GABA_B_-Rluc (0.5 μg) (as negative control) (A) and increasing amounts of cDNA for CB_2_R-GFP^2^ (0.05 to 0.6 μg) (A-C). (B, C) Cells were treated with the indicated compounds **3** (PM369), **2**, **4**, **5**, and **1** (100 nM) (see Figure 1). (D) Maximal BRET^2^ values obtained in HEK-293T cells transfected with a constant amount of cDNA for CB_2_R (WT/Val113^3.32^Met/ Val36^1.35^Met)-Rluc (0.3 μg) and increasing amounts of cDNA for CB_2_R (WT/Val113^3.32^Met/ Val36^1.35^Met)-GFP^2^ (0.05 to 0.6 μg) in the presence of compounds **2** and **3** (PM369).

To further test that the increase in CB_2_R homodimerization is triggered by the homobivalent ligand **3** (PM369), we performed similar experiments in mutant CB_2_Rs. We mutated Val113^3.32^, which is centrally located in the orthosteric cavity, and Val36^1.35^, delimiting the channel between TMs 1 and 7, to the much larger Met side chain (Figure 2D). Clearly, in the Val113^3.32^Met mutant, **3** (PM369) does not increase BRET signal, indicating that the first pharmacophore must bind the orthosteric site; and the Val36^1.35^Met mutants does not increase BRET signal either, indicating that the second pharmacophore must bind the membrane-facing pocket between TMs 1 and 7 for CB_2_R homodimerization.

In situ proximity ligation assay (PLA) is a powerful technique to detect protein-protein interactions in native tissues by assessing proximity (< 40nm) between GPCR protomers (see Methods). CB_2_R-CB_2_R homodimers were observed as red punctate staining, and the number of red spots per stained cell was significantly higher in the presence of the homobivalent ligand **3** (PM369) than in control or in the presence of the monovalent compound **2** (negative control) (Figure 3). Thus, these PLA experiments further support that the number of CB_2_R-CB_2_R homodimers are increased in the presence of PM369.

**Figure 3.**
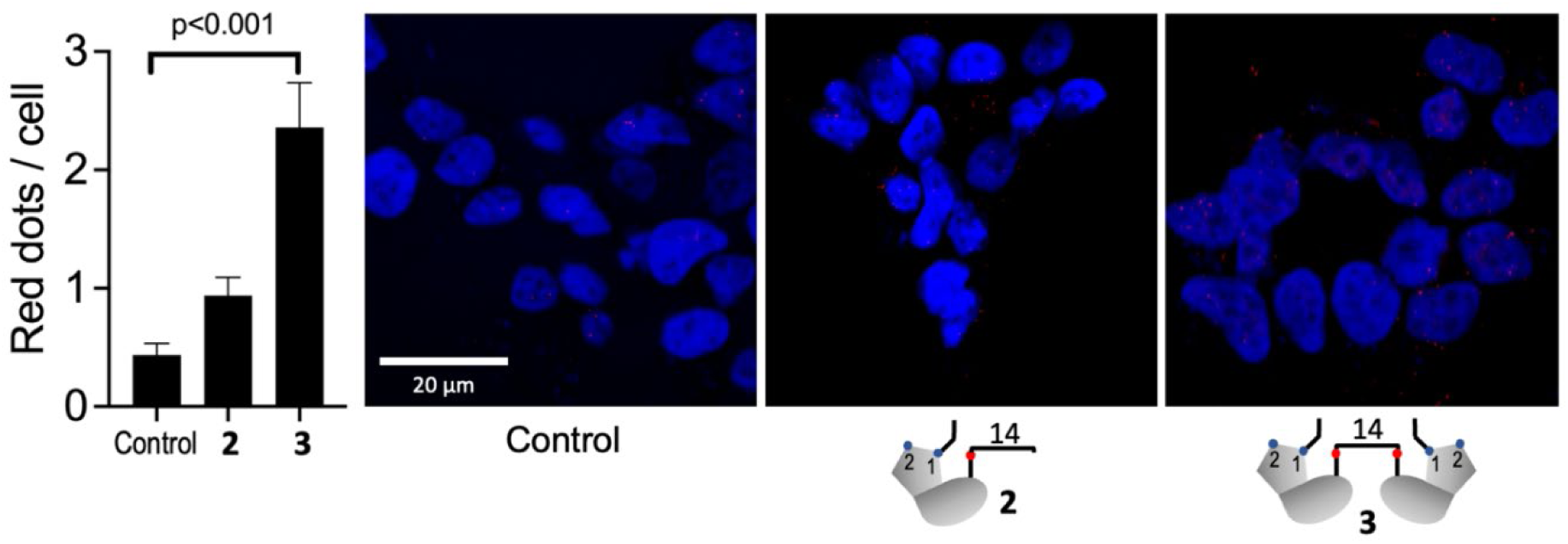
Effect of 3 (PM369) and 2 on the expression of CB_2_R-CB_2_R homomers in transfected HEK-293T cells. CB_2_R-CB_2_R heteromer expression was determined in HEK-293T cells transfected with CB_2_R-Rluc and CB_2_RGFP^2^ (0.5 µg cDNA each) by proximity ligation assay (PLA). Confocal microscopy images (superimposed sections) in which receptor complexes appear as red spots over cell stained nuclei with Hoechst (blue) were obtained in a Zeiss 880 confocal microscope. The bar graphs represent the number of red spots/cell containing spots. Values are the mean ± S.E.M. of 7 independent experiments performed in triplicates. One-way ANOVA followed by Bonferroni’s multiple comparison post hoc test was used for statistical analysis versus the control condition.

### 3.2. Computational model of homobivalent ligand PM369 bound to the CB_2_R-CB_2_R homodimer in complex with G_i_

An inactive CB_2_R protomer was added (see Methods) to our previously reported model of the active CB_2_R-G_i_ complex bound to the homobivalent N1-ethyl ligand PM369 with 14 methylene units [40]. We have recently shown that the binding pathway to the orthosteric site of CB_2_R consists of the ligand diffusing in the bilayer leaflet to contact a membrane-facing pocket between TMs 1 and 7 that is transiently occupied before entering to the orthosteric binding site through a tunnel formed between TMs 1 and 7 [39]. This membrane-facing binding site of the receptor has probably been evolutionary designed to transiently recognize endogenous cannabinoid that possess long hydrophobic moieties. It is, thus, reasonable to assume, as a working hypothesis, that the second pharmacophore of the bivalent ligand is also recognized by this favorable membrane-facing site between TMs 1 and 7 of the second protomer. Accordingly, in the constructed model (Figure 4A), the first chromenopyrazole pharmacophore of homobivalent ligand PM369 binds the orthosteric site of active CB_2_R bound to G_i_, the 14 methylene units of the spacer expand through the tunnel between TMs 1 and 7 of the active CB_2_R-G_i_ protomer, and the second chromenopyrazole pharmacophore binds the membrane-facing pocket between TMs 1 and 7 of the inactive second CB_2_R protomer. To evaluate the stability of PM369 in the CB_2_R-CB_2_R homodimer, we performed unbiased MD simulations (see Methods). Clearly, root-mean square deviations (rmsds) of the heavy atoms of the CB_2_R-G_i_-bound first pharmacophore (∼1 Å) and the CB_2_R-bound second pharmacophore (<2 Å) remained low across three replicas of 1 μs unbiased MD simulation (Supporting Information, Figure S1), suggesting that both pharmacophores optimally interact with the orthosteric and membrane-facing sites. Likewise, the rmsd values of the extended chain of 14 methylene units are also low (∼1 Å), indicating that the spacer length is optimal to bridge both sites, as determined in BRET assays above (Figure 2C). Furthermore, the low rmsd values (<2.5 Å) of the second inactive CB_2_R protomer, comparable to the active CB_2_R protomer (<2.5 Å), suggest that the initial position modeled through the TM 1/7 interface remains unchanged during the unbiased MD simulation (Supporting Information, Figure S1). Inward shifts of TMs 1 and 2 have been described during the process of activation of the closely related CB_1_R [56]. Thus, we evaluated the influence of homobivalent ligand PM369 in the extracellular conformation of TMs 1, 2 and 7 of both the active CB_2_R-G_i_ and inactive CB_2_R protomers (Supporting Information, Figure S2). The MD simulations show that PM369 does not alter the conformation of TM 2 of either active or inactive CB_2_R with respect to apo CB_2_R. Notably, inward shifts of TMs 1 (distinctive of the activation process) and 7 are observed in the active protomer, and an outward shift of TM 1 is observed in the inactive protomer. It has been previously shown that the outward conformation of TM 1, driven in this case by an antagonist of A_2A_R, facilitated the formation of the TM 1/7 interface between protomers in the A_2A_R-CB_2_R heteromer [57]. Figure 4B shows heatmaps of the predicted interactions of the second chromenopyrazole pharmacophore of PM369 with amino acids of the membrane-facing pocket between TMs 1 and 7 of the inactive CB_2_R protomer. A chemical signature of cannabinoid agonists is branched dimethyl moieties that induce receptor activation through hydrophobic interactions with Phe117^3.36^ in the *g+* conformation and Val261^6.51^ [58,59]. The pharmacophore group of homobivalent ligand PM369 contains these branched dimethyl moieties at the chromenopyrazole ring and the heptyl chain close to the aromatic ring (Figure 1). Notably, the membrane-facing site between TMs 1 and 7 contains site 1, delineated by Val36^1.35^ and Ala282^7.36^, to accommodate the dimethyl group of the chromenopyrazole ring and site 2, delineated by Leu39^1.38^ and Met286^7.40^, to accommodate the dimethyl group of the heptyl chain (Figure 4B). The highly polarizable sulfur atom of Met can form stronger hydrophobic interactions with methyl groups that aromatic or hydrophobic side chains [60]. Furthermore, this membrane-facing site contains the aromatic Phe283^7.37^ side chain that forms aromatic-aromatic interactions with the aromatic chromenopyrazole ring of the ligand, and the polar Gln32^1.31^ and Lys279^7.33^ that act as hydrogen bond donor in the interaction with the N2 atom of the N1-ethyl isomer of the pyrazole ring. Importantly, the N2-ethyl isomer cannot achieve this key interaction, due to a different orientation of the N1 atom, which explains that homobivalent ligand **5** could not increase BRET values (Figure 2B).

**Figure 4.**
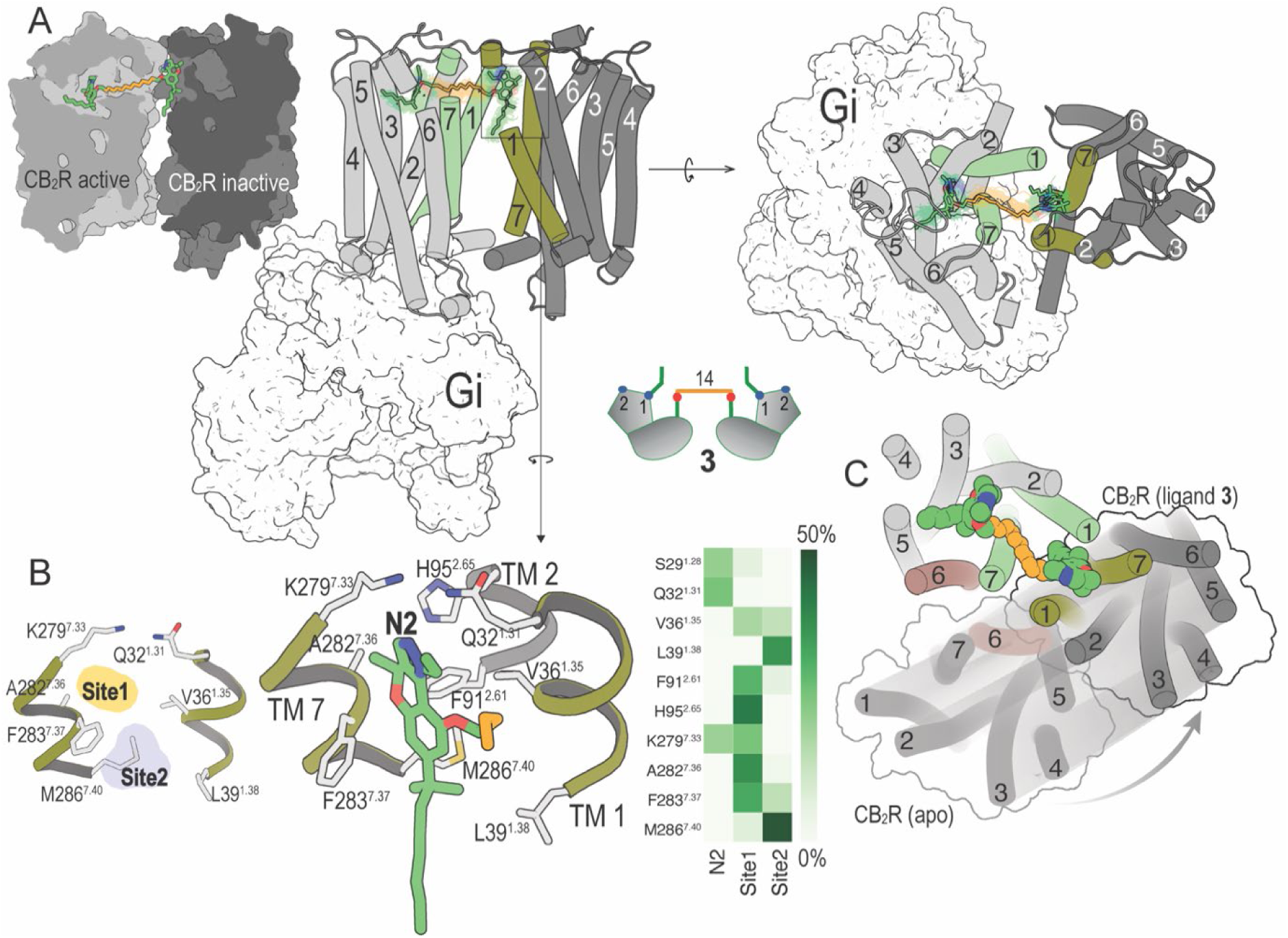
Computational model of the CB_2_R-CB_2_R homodimer in complex with G_i_. (A) Cross-section through the CB_2_R-CB_2_R homodimer, highlighting homobivalent ligand **3** (PM369, color sticks) binding the orthosteric site of active CB_2_R and the membrane-facing pocket of inactive CB_2_R. A representative structure (color sticks) and 100 structures collected every 10 ns (translucent lines) of homobivalent ligand **3** (PM369), as devised from unbiased 1μs MD simulations in views parallel and perpendicular to the membrane plane. The structure of CB_2_R-CB_2_R (TM helices are depicted as cylinders) in complex with G_i_ (white surface) corresponds to the initial structure. One replica is displayed for better visualization although additional replicas showed consistent behavior (Supporting Information, Figure S1). (B) Detailed views and heatmaps (calculated with GetContacts, https://getcontacts.github.io/interactions.html) depicting the predicted interactions between the aromatic chromenopyrazole ring of **3** (PM369) and Phe283^7.37^ of inactive CB_2_R protomer, between the N2 atom of the N1-ethyl isomer of the pyrazole ring and Gln32^1.31^ and Lys279^7.33^, and between branched dimethyl moieties at the chromenopyrazole ring and the heptyl chain with Val36^1.35^ and Ala282^7.36^ (site 1), and Leu39^1.38^ and Met286^7.40^ (site 2), respectively. (C) Computational molecular models of the CB_2_R-CB_2_R homodimer built using the TM 6 interface (in red) for apo (previously determined by BiFC assays in combination with disruptive peptides [57] and the TM 1/7 interface (in green) for homobivalent ligand PM369. To facilitate visualization of the dynamic dimerization interface of the CB_2_R-CB_2_R homodimer, the TM 6 and TM 1/7 interfaces are displayed in the same panel, as well as the geometrical morphing between both conformations.

### 3.3. Homobivalent ligand PM369 potentiates cellular signaling

We have first compared the agonist-induced response of the chromenopyrazole pharmacophore **1** with JWH-133, a potent and selective CB_2_R agonist, both binding exclusively at the orthosteric site. Cells stimulated with forskolin and treated with JWH-133 or compound **1** showed reduced cAMP production, as expected for G_i_-coupled receptors (Figure 5A). However, the chromenopyrazole **1** is considered a partial agonist referred to JWH-133 due to a decrease of cAMP of less magnitude. Notably, homobivalent ligand PM369 is as efficient as JWH-133 in reducing cAMP production unlike its corresponding monovalent ligand **2** that lacks the second pharmacophore (negative control). Moreover, PM369 left-shifts the dose-response curve by one log unit relative to JWH-133, indicating an increase in potency.

**Figure 5.**
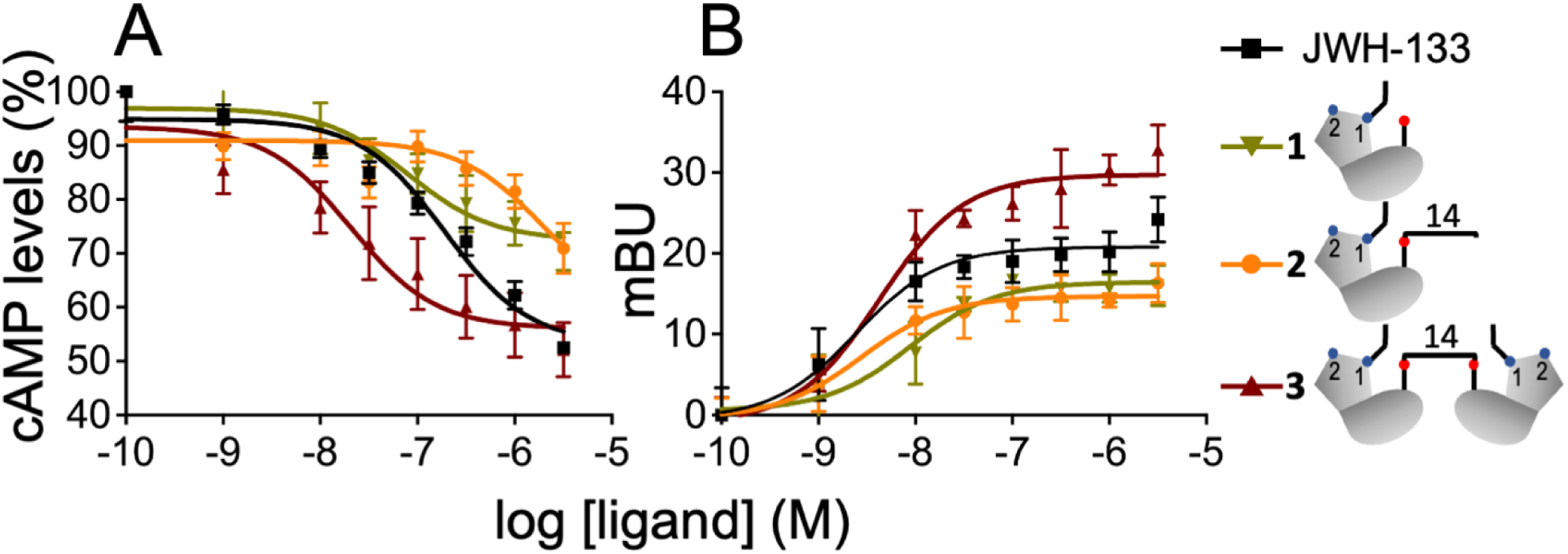
Functional characterization of the CB_2_R-CB_2_R homodimer in transfected HEK-293T cells. (A) cAMP levels were determined in HEK-293T cells transfected with CB_2_R (0.5 μg cDNA). (B) β-arrestin 2 recruitment experiments were performed in HEK-293T cells transfected with β-arrestin 2-Rluc (0.5 μg) and CB_2_R-YFP (0.5 μg) cDNAs. (A, B) Cells were stimulated with increasing concentrations (0.1 NM to 3 µM) of the full agonist JWH-133 (100 nM) or with compounds **1**, **2**, and **3** (PM369). cAMP accumulation was collected upon stimulation with forskolin (Fk, 500 nM).

We aim to understand the mechanisms by which the second pharmacophore of PM369 that binds to the membrane-facing pocket between TMs 1 and 7 of the inactive second protomer enhances agonist-induced receptor activation. Previous work using bimolecular fluorescence complementation (BiFC) experiments, in which HEK-293T cells were co-transfected with CB_2_Rs separately fused to complementary halves of the yellow fluorescent protein (N-terminal, CB_2_R-nYFP; or C-terminal, CB_2_R-cYFP), in combination with disruptive peptides targeting a specific dimer interface, showed that fluorescence of CB_2_R-nYFP/CB_2_R-cYFP was mainly reduced by the TM6 peptide of CB_2_R with minor, but statistically significant, contributions of TM1, TM5, and TM7 [57]. This suggested that, in the absence of the ligand (apo CB_2_R), the CB_2_R homodimer is dynamically tuneable, and several dimer interfaces co-exist and interconvert as also described for neurotensin receptor 1 [13]. In a CB_2_R-CB_2_R homodimer formed through the TM 6 interface, one of the protomers might partially prevent the outward movement of TM 6 of the partner protomer, challenging the opening of the intracellular cavity for receptor activation and G_i_ binding. Therefore, homobivalent ligand PM369 has two effects in the process of receptor activation. First, binding of the second pharmacophore group to the membrane-facing pocket between TMs 1 and 7 of the inactive second protomer interconverts the CB_2_R-CB_2_R homodimer from TM 6 to the TM 1/7 interface, facilitating the opening of the intracellular cavity for G protein binding at the partner receptor. Second, binding of the first pharmacophore group to the orthosteric site activates the molecular switches for receptor activation with improved efficacy, like the full agonist JWH-133. The increase in potency is attributed to the multiple contacts of the second pharmacophore with the membrane-facing site between TMs 1 and 7 (Figure 4B). Figure 4C shows the proposed structure of the CB_2_R-CB_2_R homodimer bound to the homobivalent ligand PM369, together with apo CB_2_R-CB_2_R homodimer, constructed via the TM 6 interface, to facilitate visualization of the dynamic dimerization interfaces of the homodimer.

The intracellular cavity, opened upon receptor activation by the outward movement of TM 6 to accommodate the C-terminal α5 helix of the Gα subunit [47], also binds the finger loop of arrestin [50]. Accordingly, homobivalent ligand PM369 should also have a significant effect in β-arrestin-2 recruitment. Like in cAMP measurements, JWH-133 recruited β-arrestin-2 more efficiently than the chromenopyrazole pharmacophore **1** (Figure 5B). Surprisingly, PM369 not only improved β-arrestin-2 recruitment relative to chromenopyrazole **1**, but also surpassed the efficacy of the full agonist JWH-133, unlike the corresponding monovalent ligand **2** (negative control). Therefore, the TM 1/7 interface of the CB_2_R-CB_2_R homodimer, triggered by PM369, facilitates β-arrestin-2 recruitment.

### 3.4. Computational model of the CB_2_R-CB_2_R homodimer in complex with β-arrestin

To understand the mechanism by which homodimerization via the TM 1/7 interface facilitates β-arrestin-2 recruitment, we constructed a computational model of the CB_2_R-CB_2_R homodimer bound to β-arrestin-2 (see Methods). Arrestins are organized in a fold consisting of N- and C-domains [61], at whose interface is the finger loop that inserts into the receptor intracellular binding cavity opened with the outward movement of TMs 5 and 6 (Figure 6A). Biophysical and structural studies have revealed conformational variability of arrestin while coupled to the receptor [62]. These multiple conformations are shown in Figure 6B in which the available GPCR-bound β-arrestin structures are superimposed to the active protomer of the CB_2_R-CB_2_R homodimer. Two clusters of arrestin orientations can be visually distinguished: the NTS_1_R-bound and the V_2_R/5-HT_2B_R/β_1_R/CB_1_R^(8WU1)^/M_2_R/CB_1_R^(8WRZ)^- bound orientations. Furthermore, a lateral view of the structures forming the later cluster also shows two distinct tilt angles of β-arrestin relative to the bound receptor (Figures 6C-6D). However, two different groups have recently published the structure of β-arrestin bound to the highly homologous CB_1_R. Liao et al. [50] showed that the tilt angle of β-arrestin bound to CB_1_R (72°) is like the complexes formed with β_1_R (75°) or the muscarinic M2 receptor (75°) (Figure 6C). In contrast, the structure reported by Wang et al. [51] adopts a tilt angle of 46° comparable to the value observed in the complex with V_2_R (Figure 6D). In both structures the C-domain of β-arrestin is oriented towards the cell membrane, where the C-edge loop acts as a membrane anchor [59]; whereas Lys158 localized at the tip of the N-domain of β-arrestin is facing the inactive protomer of the CB_2_R-CB_2_R homodimer (Figures 6E-6F). However, in the 8WRZ structure of Wang et al. Lys158 points towards ICL3 of the inactive protomer (Figure 6F), whereas in the 8WU1 structure of Liao et al. Lys158 points towards H8 and it is Arg162 that points towards ICL3 (Figure 6E). Unfortunately, the interactions of Lys158 with the inactive CB_2_R protomer cannot be assigned non-arbitrarily, as they depend on the modeled conformation of β-arrestin. However, this analysis supports our proposal that β-arrestin stablishes new contacts with the inactive CB_2_R protomer of the CB_2_R-CB_2_R homodimer in the TM 1/7 interface.

**Figure 6.**
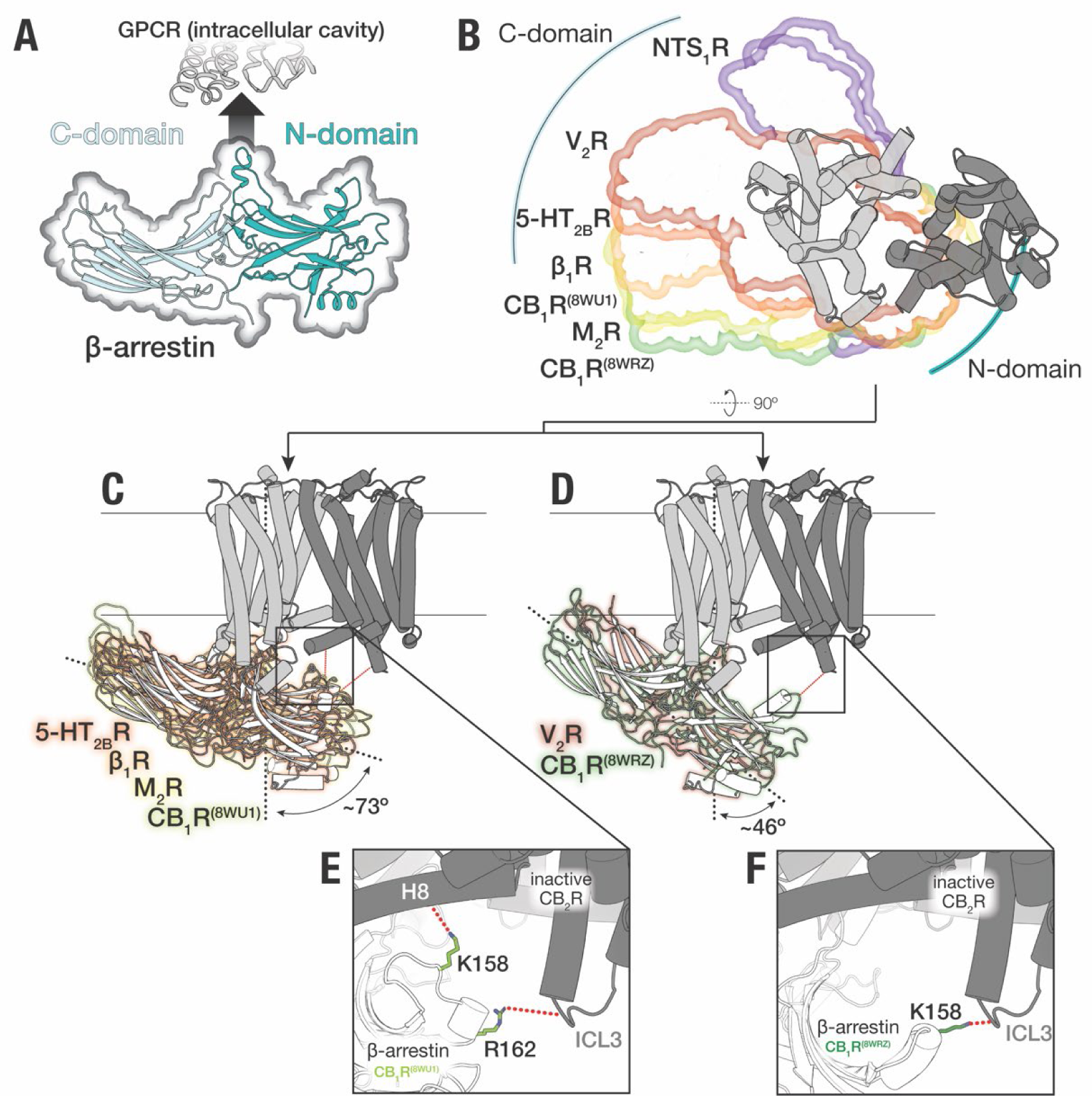
Computational model of the CB_2_R-CB_2_R homodimer in complex with β-arrestin 2. (A) Structure of β-arrestin in a fold consisting of C- and N-domains, in light cyan and teal, respectively, at whose interface is the finger loop that inserts into the receptor intracellular binding cavity. (B) Available GPCR-bound β-arrestin structures [NTS_1_R- (6UP7, 6PWC)/V_2_R- (7R0C)/5-HT_2B_R- (7SRS)/β_1_R- (6TKO)/M_2_R-(6U1N)/CB_1_R- (8WU1, 8WRZ) bound) superimposed to the active protomer (light gray) of the CB_2_R-CB_2_R homodimer. (C, D) Lateral view of GPCR-bound β-arrestin structures showing two distinct tilt angles of β-arrestin relative to the bound receptor. (E, F) Detailed views of the interaction between Lys158 located at the tip of the N-domain of β-arrestin and the inactive CB_2_R protomer.

## 4. Conclusion

Bitopic and bivalent ligands are single chemical entities composed of two pharmacophore units, covalently linked by an appropriate spacer, that bind the orthosteric site as well as a less conserved site within the same receptor unit or bridge two receptor units [27][28]. Many of the synthesized bivalent ligands for cannabinoid receptors in particular, and GPCRs in general, contain spacers too short for simultaneous binding of both orthosteric sites of the (homo/hetero)dimer [63][64]. It has been suggested that these short bivalent ligands might bind the orthosteric pocket of one protomer and a complementary site, different from the orthosteric, of the second protomer [64]. Alternatively, these short ligands can bind to both orthosteric sites of the (homo/hetero)dimer, not simultaneously, but via a ‘flip-flop’ mechanism or cooperatively [65].

Here, we have tested whether previously reported compounds by us, formed by equal chromenopyrazole moieties as pharmacophores connected by spacers whose lengths vary from six to sixteen methylene units [40], can modulate the dynamics of CB_2_R homodimerization by simultaneously binding both protomers of the CB_2_R-CB_2_R homodimer. Notably, only homobivalent ligand PM369 with 14 methylene units, but none of the other tested compounds, increased BRET values relative to untreated cells. Computational and experimental results showed that the first chromenopyrazole pharmacophore of PM369 binds the orthosteric site of active CB_2_R bound to G_i_, the 14 methylene units of the spacer expand through the tunnel between TMs 1 and 7 of the active CB_2_R-G_i_ protomer, and the second chromenopyrazole pharmacophore binds the membrane-facing pocket between TMs 1 and 7 (complementary site) of the inactive second CB_2_R protomer. This binding mode of PM369 triggers the formation of the CB_2_R-CB_2_R homodimer via the TM 1/7 interface. This provides unique pharmacological properties, such as increased potency in G_i_ binding and increased recruitment of β-arrestin. It was previously shown by others that oligomerization of the platelet-activating factor receptor either via TM 1 or TM 4/5 interfaces decreased recruitment of β-arrestin [22]. Therefore, these results suggest that only certain conformation(s) of the homodimer, stabilized in this work by homobivalent ligand PM369, is(are) able to recruit β-arrestin more efficiently.

By promoting homodimerization and altering downstream signaling events, PM369 is explored as a tool to better understand GPCR pharmacology. However, these findings also suggest that PM369 could offer new therapeutic strategies by controlling the signaling pathways mediated by CB_2_R. CB_2_R plays a crucial role in modulating immune responses and inflammation [66]. Its upregulation under pathological conditions and its association with anti-inflammatory effects highlight the therapeutic potential of targeting CB_2_R in various disease states such as autoimmune diseases, neuroinflammation, and tissue injury [37,38].

Altogether, these novel results open new possibilities to control GPCR signaling.

## Credit authorship contribution statement

Gemma Brugal, Paula Morales, and Marc Gómez-Autet contributed equally to this work. Gemma Brugal: Investigation. Methodology in pharmacology. Data curation.

Marc Gómez-Autet: Investigation. Methodology in molecular dynamics. Data curation. Paula Morales: Investigation. Methodology in medicinal chemistry. Data curation.

Joan Biel Rebassa: Investigation. Methodology in pharmacology. Data curation.

Claudia Llinas del Torrent: Investigation. Methodology in molecular dynamics. Data curation.

Nadine Jagerovic: Medicinal chemistry conceptualization. Funding acquisition. Writing, review & editing. Co-corresponding author.

Leonardo Pardo: Molecular dynamics conceptualization. Funding acquisition. Writing, review & editing. Co-corresponding author.

Rafael Franco: Pharmacological conceptualization. Funding acquisition. Writing, review & editing. Co-corresponding author.

All authors read and approved the final manuscript for publication.

## Data availability

Data will be made available on request.

## Supporting information

Supplementary data associated with this article can be found in the online version at

## ABBREVIATIONS

CB_1_R: cannabinoid receptors type 1 (CB_1_R)
CB_2_R: cannabinoid receptors type 2
GPCR: G protein-coupled receptor
cAMP: cyclic adenosine monophosphate
TM: transmembrane helix
MD: molecular dynamics
BRET: bioluminescence resonance energy transfer
Rluc: Renilla luciferase
PLA: proximity ligation assay
GFP: green fluorescent protein
Rmsd: root-mean square deviation
BiFC: bimolecular fluorescence complementation
YFP: yellow fluorescent protein

