## Supplemental Figures S1 and S2 for "Homodimerization of CB_2_ cannabinoid receptor triggered by a bivalent ligand enhances cellular signaling"

[a]Department of Biochemistry and Physiology. Faculty of Pharmacy and Food Sciences. Universitat de Barcelona. 08028 Barcelona. Spain

[b]Institute of Neuroscience, University of Barcelona (NeuroUB), 08035 Barcelona, Spain

[c]Centro de Investigación en Red, Enfermedades Neurodegenerativas (CIBERNED), Instituto de Salud Carlos III, 28031 Madrid

[d]Laboratory of Computational Medicine, Biostatistics Unit, Faculty of Medicine, Universitat Autònoma de Barcelona, 08193 Bellaterra, Spain

[e]Medicinal Chemistry Institute, Spanish National Research Council, 28006 Madrid, Spain

[f]Department of Biochemistry and Molecular Biomedicine, Faculty of Biology, Universitat de Barcelona, 08028 Barcelona, Spain

[π]These authors contributed equally to this work

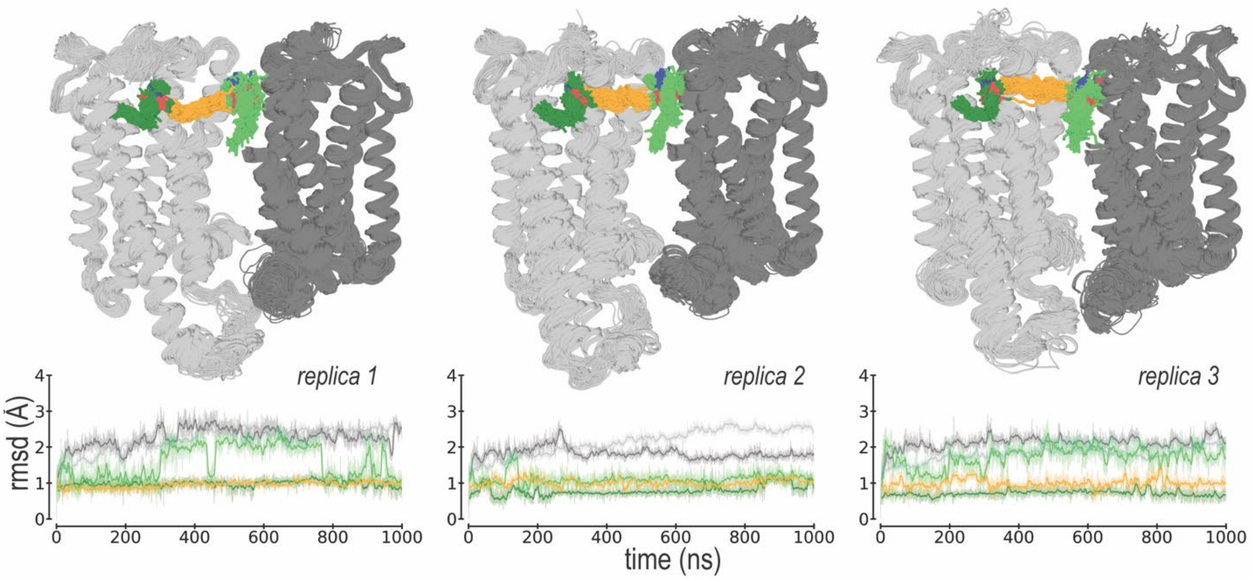


**Figure S1**. **Molecular dynamic simulations of the $CB_2R$-$CB_2R$ homodimer in complex with $G_i$**. Evolution of the first chromenopyrazole pharmacophore (dark green), binding to the orthosteric site of active $CB_2R$ (light gray) bound to $G_i$ (not shown for clarity), the 14 methylene units of the spacer (orange), and the second chromenopyrazole pharmacophore (light green), binding to the membrane-facing pocket of inactive $CB_2R$ (dark gray), of homobivalent ligand PM369 as devised from three replicas of unbiased 1$\mu$s MD simulations. The stabilities of these moieties of PM369 were analyzed via root mean-square deviations (rmsd) of the heavy atoms, and the stabilities of active and inactive $CB_2R$ were analyzed via rmsd of the backbone atoms.

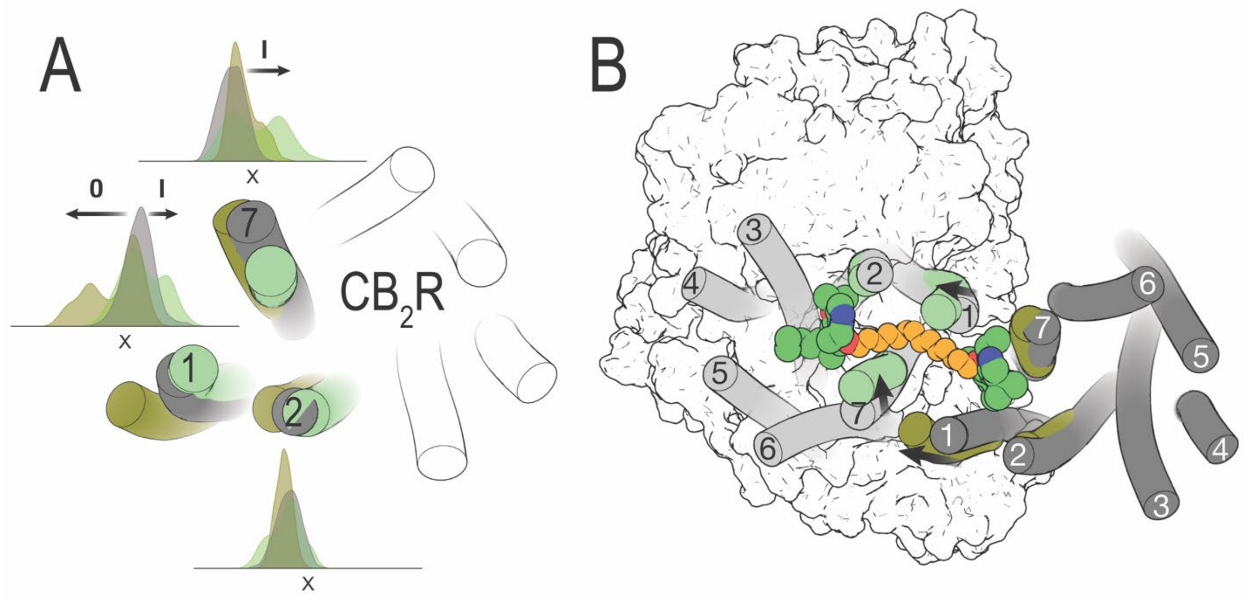


**Figure S2. Homobivalent ligand PM369 driven rearrangements of TMs 1, 2 and 7.** (A) Superimposition of the conformation of TMs 1, 2 and 7 as observed in MD simulations of apo $CB_2R$ (grey), and inactive $CB_2R$ (olive) and active $CB_2R$-$G_i$ (light green) as observed in MD simulations of homobivalent ligand PM369 bound to the $CB_2R$-$CB_2R$ homodimer in complex with $G_i$. Distribution of the x-values of the center of mass of the amino acids 1.32-1.35 in TM 1, 2.52-2.55 in TM 2, and 7.31-7.34 in TM 7 of $CB_2R$ during three replicas of unbiased 1 µs MD simulation are shown. Arrows show the most remarkable Inward and Outward movements. The xy plane is as defined by the Orientations of Proteins in Membranes (OPM). (B) Significant movements (black arrows) of TMs 1 and 7 observed in the MD simulations of homobivalent ligand PM369 bound to the $CB_2R$-$CB_2R$ homodimer in complex with $G_i$, relative to their orientation in the initial structure (grey).
